# Somatic mutations distinguish melanocyte subpopulations in human skin

**DOI:** 10.1101/2025.02.07.637114

**Authors:** Bishal Tandukar, Delahny Deivendran, Limin Chen, Neda Bahrani, Beatrice Weier, Harsh Sharma, Noel Cruz-Pacheco, Min Hu, Kayla Marks, Rebecca G Zitnay, Aravind K. Bandari, Rojina Nekoonam, Iwei Yeh, Robert Judson-Torres, A. Hunter Shain

## Abstract

To better understand the homeostatic mechanisms governing melanocytes, we performed deep phenotyping of clonal expansions of single melanocytes from human skin. In total, we interrogated the mutational landscapes, gene expression profiles, and morphological features of 297 melanocytes from 31 donors. To our surprise, a population of melanocytes with low mutation burden was maintained in sun damaged skin. These melanocytes were more stem-like, smaller, less dendritic and displayed distinct gene expression profiles compared to their counterparts with high mutation burdens. We used single-cell spatial transcriptomics (10X Xenium) to reveal the spatial distribution of melanocytes inferred to have low and high mutation burdens (LowMut and HighMut cells), based on their gene expression profiles. LowMut melanocytes were found in hair follicles as well as in the interfollicular epidermis, whereas HighMut melanocytes resided almost exclusively in the interfollicular epidermis. We propose that melanocytes in the hair follicle occupy a privileged niche, protected from UV radiation, but periodically migrate out of the hair follicle to replenish the interfollicular epidermis after waves of photodamage. More broadly, our study illustrates the advantages of a cell atlas that includes mutational information, as cells can change their cellular states and positional coordinates over time, but mutations are like scars, providing a historical record of the homeostatic processes that were operative on each cell.

## Introduction

During embryogenesis, neural crest cells give rise to smooth muscle, bone, connective tissue, neurons, and melanocytes^1–3^. Melanocytes are pigment producing cells that mostly migrate to the skin but can colonize, to a lesser extent, other sites in the body. The developmental trajectory of the melanocytic lineage has predominantly been traced in animal models^1–3^, but there are key differences between melanocytes from humans and other animals. Fish, birds, and rodents evolved pigmentation mainly to alter their appearance (e.g. for camouflage or mating purposes)^4^, whereas in humans, pigmentation evolved as a defense mechanism against UV radiation^5^.

Single-cell RNA-sequencing has helped define the cellular fates of melanocytes in humans^6^. As an example, melanocytes from acral body sites (the non-hair bearing skin of the foot sole, palm, and nail bed) occupy a distinct cellular state from the cutaneous melanocytes found elsewhere in the body^6^, likely explaining why acral melanomas are biologically different from cutaneous melanomas^7,8^. While RNA-sequencing has illuminated anatomic specificity within melanocytes in humans, our understanding of their cellular hierarchies remains incompletely understood.

To address this gap in knowledge, we catalogued the mutational, gene expression, and morphological features of small colonies of individual melanocytes expanded *ex vivo* from human skin (Fig. S1A). Cell atlas studies are routinely carried out by performing single-cell RNA sequencing to define cellular states^9^. Our multi-omic approach has advantages over single-cell RNA-sequencing alone (Fig. S1B). Single-cell RNA-sequencing provides a one-dimensional view into the biological state of each cell, and the shallow sequencing depth from these technologies imparts limited resolution into heterogeneity within a cellular lineage. Our work revealed mutationally distinct subpopulations of melanocytes in the skin. Moreover, we validated their existence and interrogated their spatial distribution with spatial transcriptomics (Xenium Analyzer, 10X Genomics).

## Results

### Phenotyping melanocytes from human skin

Melanocytes were collected from punch biopsies of sun-exposed adult skin. After incubation in dispase to break down extracellular matrix proteins, the epidermis was physically separated from the dermis with tweezers. Bulk melanocytes were established in tissue culture from the epidermal compartment as described (see methods). This separation strategy is thought to enrich for melanocytes from the interfollicular epidermis^10^.

We catalogued somatic mutations in melanocytes by clonally expanding individual melanocytes into small colonies (∼230 cells), further amplifying the DNA and RNA of each colony *in vitro*, and sequencing the amplified nucleic acids (Fig. S1A). We developed a bioinformatic approach, which we previously described^11^, to call mutations at high sensitivity and specificity from each expansion of melanocytes. Since we sequenced both DNA and RNA, there was matching gene expression data to accompany the mutational information from each colony. The RNA was sequenced at higher depth than a typical cell atlas study^12,13^, providing higher resolution into lineage subtypes. Furthermore, we photographed each colony to record their morphological features.

A limitation to our approach is the involvement of cell culture, which may alter the gene expression, morphological, or, to a lesser extent, the mutational state of each cell. We also characterized fewer cells than a typical cell atlas study. However, these limitations are counterbalanced by the extensive phenotypic information that was collected from each cell’s colony (Fig. S1B). In total, we profiled 297 cells from 58 independent skin biopsies of 31 unique donors – approximately half were previously published^11^ and the remaining are newly sequenced here.

### Low- and high-mutation burden melanocytes co-exist in human skin

The mutation burden of individual melanocytes varied from person to person, site to site within each person, and cell to cell within each site^11^. In this study, we focused on the latter source of variability, aiming to understand how melanocytes from the same skin biopsy, which would be expected to experience similar exposures to UV radiation, accumulate different levels of mutational damage.

Towards this goal, we plotted the mutation burdens of individual melanocytes versus the mutation burdens of the tissues from which they were derived (Fig. 1). Skin samples with high mutation burdens, on average, maintained a subpopulation of melanocytes with low mutation burdens. We defined melanocytes with low- and high-mutation burdens as those in the bottom and top quartile of mutation burdens. We also restricted our comparisons to melanocytes from tissues whose mutation burden exceeded 3 mutations/Megabase (mut/Mb) – this decision was due to the fact that it was difficult to separate low- and high-mutation burden melanocytes when there was little mutational damage in the tissue. The gates defining the high- and low-mutation burden melanocytes are shown in figure 1.

**Figure 1.**
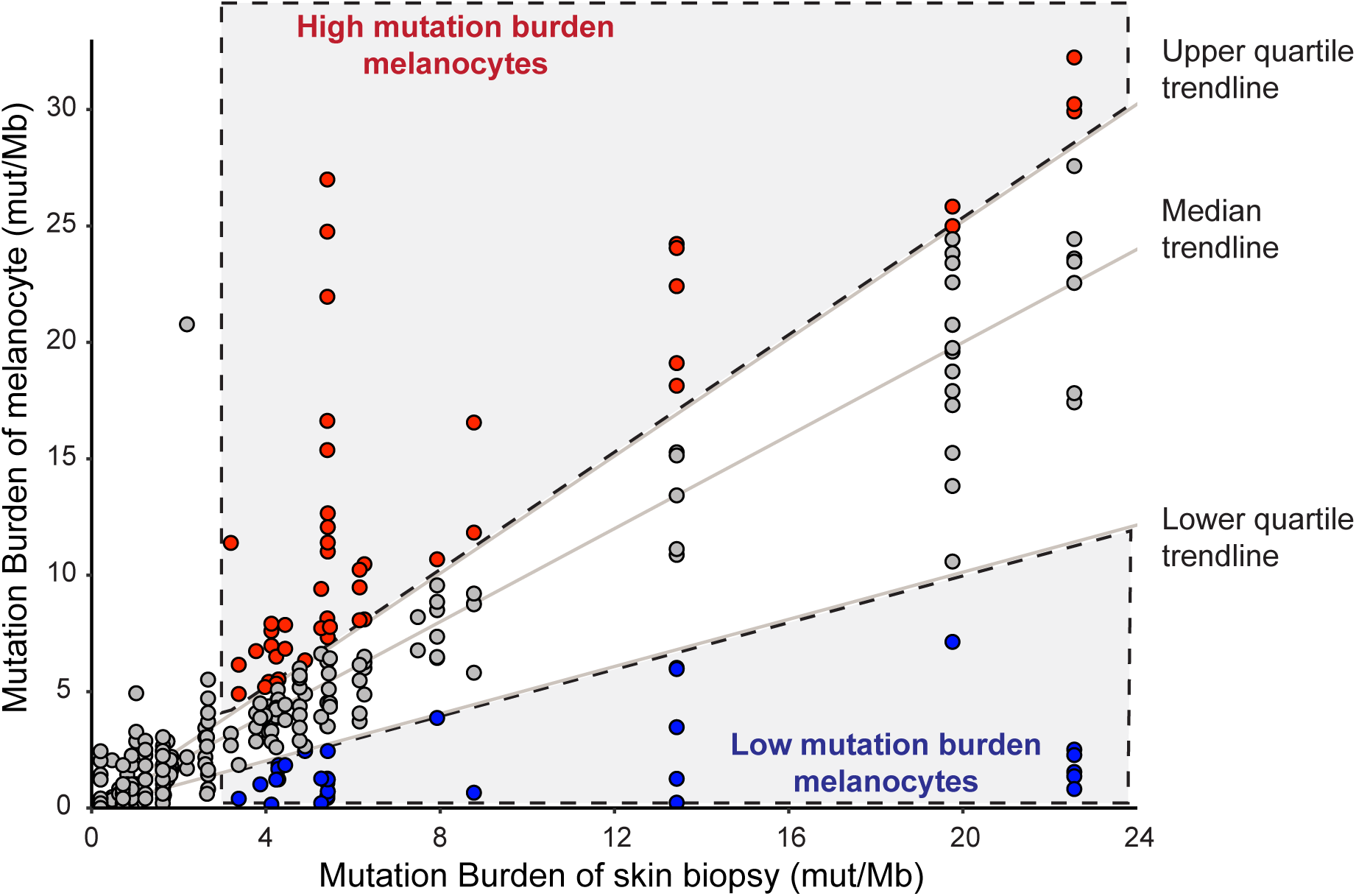
Melanocytes with high- and low-mutation burdens coexist in human skin biopsies. The mutation burdens (mutations/megabase, mut/Mb) of individual melanocytes are plotted versus the mutation burdens of the skin biopsies from which they were collected. The mutation burden of each skin biopsy is defined as the median mutation burden of all melanocytes from that biopsy. Upper, middle, and lower quartile trendlines are shown. Low- and high-mutation burden melanocytes were gated (see methods) as shown. Note how skin samples with high mutation burdens maintain populations of melanocytes with few mutations.

### Distinct mutational factors operate on melanocytes with high and low mutation burdens

Next, we characterized the types of mutations in the two populations of melanocytes. The melanocytes with high mutation burdens had a greater proportion of cytosine to thymine transitions at the 3’ basepair of dipyrimidines (Fig. 2) – the classic mutation associated with UV radiation^14^. They also had a greater proportion of signature 7 mutations (Fig. 2D) – a signature defined in pan-cancer analyses that is attributable to UV radiation^15^. By contrast, the low mutation burden melanocytes had a greater proportion of “clock-like” signatures 1 and 5^15^ (Fig. 2E). These observations imply there is a subpopulation of melanocytes in heavily sun exposed skin that remains relatively protected from UV radiation.

**Figure 2.**
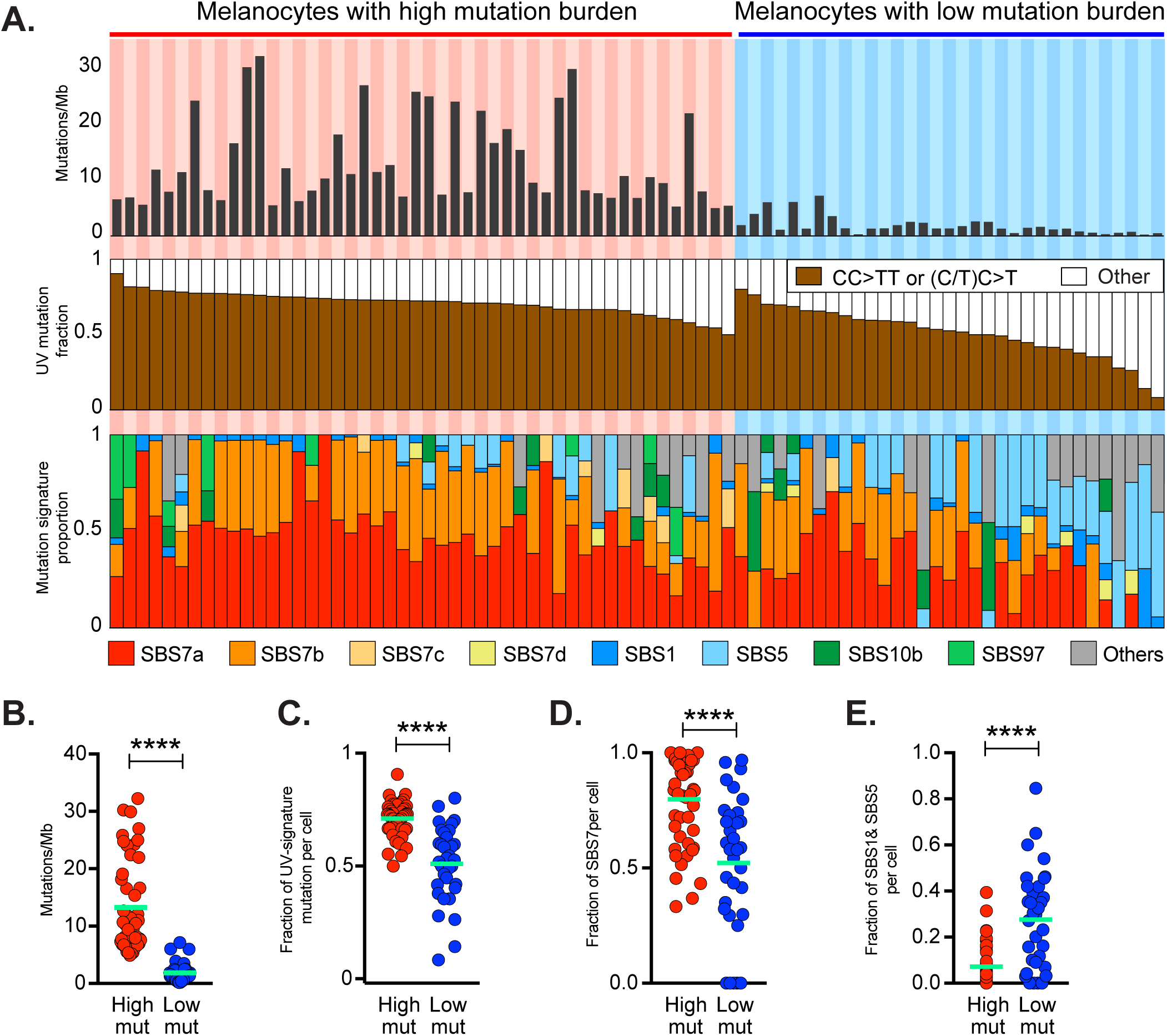
Distinct mutational factors operate on melanocytes with high- and low-mutation burdens. **A.** The top panel shows mutation burdens of site-matched melanocytes with high- and low-mutation burdens (see Fig. 1 for definition of each group). Each column represents one melanocyte. The middle panel shows the fraction of UV-radiation-induced mutations in each melanocyte, and the bottom panel shows fractions of mutational signatures in each cell. The melanocytes are arranged in descending order by their fraction of UV-induced mutations (middle panel). The dot plots summarize select features in high- and low-mutation burden melanocytes, including mutation burden **(B)**, fraction of UV-radiation-induced mutations per cell **(C)**, fraction of SBS7-associated mutations per cell **(D)**, as well as fraction of SBS1 and SBS5 associated mutations per cell **(E)**. The green bars represent the mean values for each subset, while each dot corresponds to the value for an individual melanocyte. Statistical significances are based on the Wilcoxon rank-sum test *(****p < 0.*0001 for cell-to-cell comparisons).

### Low- and high-mutation burden melanocytes have distinct gene expression profiles

To further understand the biological differences between melanocytes with low- and high-mutation burdens, we performed differential gene expression analysis (Fig. 3A). As expected, both groups expressed melanocyte markers at levels that were several orders of magnitude greater than keratinocytes or fibroblasts (Fig. 3B, upper heatmap), confirming their shared melanocytic lineage despite phenotypic variation within the lineage. The significantly up-regulated genes in melanocytes with high- and low-mutation burdens were compared to the molecular signature database^16^. All gene sets with significant overlap are listed in Table S4, and a subset of notable overlaps (discussed below) are highlighted in figure 3C.

**Figure 3.**
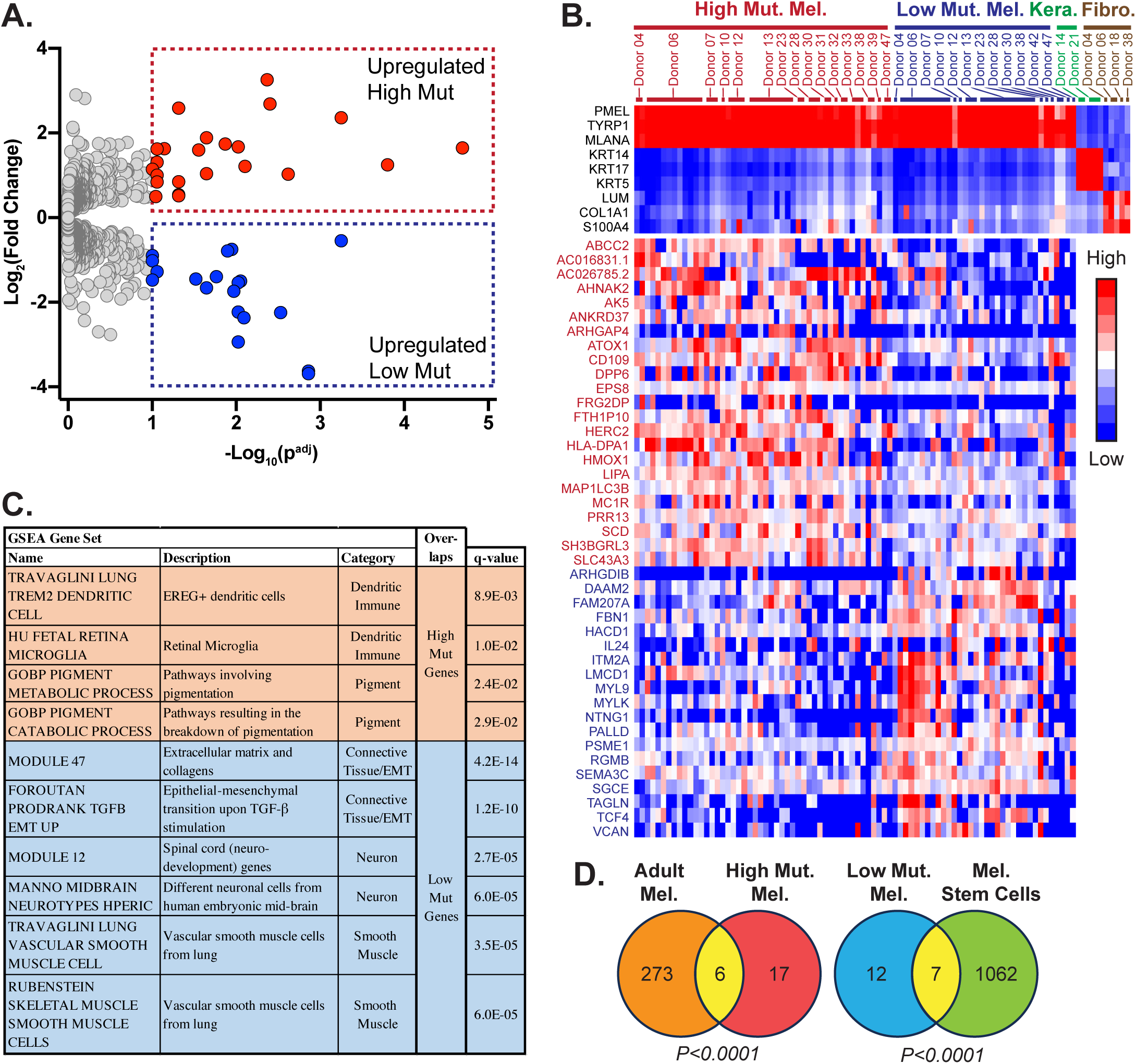
Distinct gene expression profiles in melanocytes with high and low mutation burdens. **A.** Volcano plot shows differentially expressed genes between melanocytes with high- and low-mutation burdens using DESeq2. Significant genes had adjusted p-values of less than 10% (Benjamini-Hochberg). **B.** A heatmap shows expression of genes (rows) in individual skin cells (columns). The top portion compares melanocytes to other cell types in the skin (keratinocytes and fibroblasts) using selected markers known to be expressed in each lineage. The bottom portion shows differentially expressed genes (from panel A) in melanocytes with high and low mutation burdens. **C.** Select gene sets (see table S4 for full list) from the molecular signature database that overlap with high mutation burden genes or low mutation burden genes. **D.** Venn diagram shows overlap/non-overlap between genes discovered in our study (the high and low mutation burden genes) and genes discovered from Belote *et al.* Nature Cell Biology, 2021 (“Adult Melanocyte” or “Melanocyte Stem Cell” genes). *p*-values calculated with hypergeometric test.

The melanocytes with high mutation burdens expressed higher levels of genes involved in pigmentation and antigen presentation (Fig. 3B, C). Examples include *HMOX1*, *ABCC2*, and *MC1R*, whose proteins participate in metabolism and catabolism of pigmentation products. Melanocytes with high mutation burdens also expressed *HERC2*, which has a well-established genetic linkage to skin tone, eye color, and melanoma susceptibility in genome-wide association studies^17^. Finally, they expressed genes overlapping with previously described immune signatures (Table S4 and Fig. 3C), including genes whose products are responsible for protein breakdown (*LIPA*, *HMOX1*) and antigen presentation (*HLA-DPA1*).

The melanocytes with low mutation burdens expressed higher levels of genes typically observed in cells from neuronal, smooth muscle, and connective tissues (Fig. 3C). This was notable because, during development, neural crest cells give rise to each of these cell types in addition to melanocytes. Examples of neural crest genes include: *VCAN*, *FBN1*, *PALLD*, *ITM2A* (connective tissue genes); *TAGLN*, *MYL9*, *MYLK*, *SGCE*, *HACD1* (smooth muscle genes); and *SEMA3C*, *TCF4*, *DAAM2*, *RGMB*, *NTNG1* (neuronal genes).

We also compared the genes, upregulated in low- and high-mutation burden melanocytes, to gene sets defined in cell atlas studies focused on melanocytes^6,18^ (Fig. 3D, S2). The genes upregulated in our high mutation burden melanocytes overlapped with genes upregulated in “adult” melanocytes from Belote and colleagues, and the genes upregulated in our low mutation burden melanocytes overlapped with “melanocyte stem cell (MSC)” genes from Belote and colleagues. The “MSC” gene set, from Belote *et. al.*, was enriched in melanocytes from fetal tissue and suggested to derive from melanocytes in hair follicles^6^.

Finally, we used a recently developed tool, WIMMS (What Is My Melanocytic Signature), to interrogate how our gene expression signatures relate to other melanocytic signatures (Fig. S3)^19^. Briefly, WIMMS re-interpreted 39 commonly referenced gene expression signatures from studies of melanoma or melanocytes, showing that many of the signatures are capturing overlapping cell states. With this framework, our high mutation burden melanocytes are most similar to “Differentiated” melanocytes from other studies, while our low mutation burden melanocytes are most similar to “AXL”, “Neuronal”, and “Invasive” melanocytes from other studies.

Taken together, gene expression analyses suggest that melanocytes with high mutation burdens occupy a relatively differentiated state as compared to low mutation burden melanocytes.

### Low- and high-mutation burden melanocytes have different morphological features

Next, we compared the morphology of melanocytes with high- and low-mutation burdens (see Fig. 4A-B for representative examples of each). Melanocytes that could be distinguished from their neighbors were segmented, and we calculated their perimeters, surface areas, and number of dendritic extensions. Melanocytes with high mutation burdens had more complex morphologies, defined by the ratio of their perimeter to surface area (Fig. 4C). They also tended to be dendritic and larger (Fig 4D, E).

**Figure 4.**
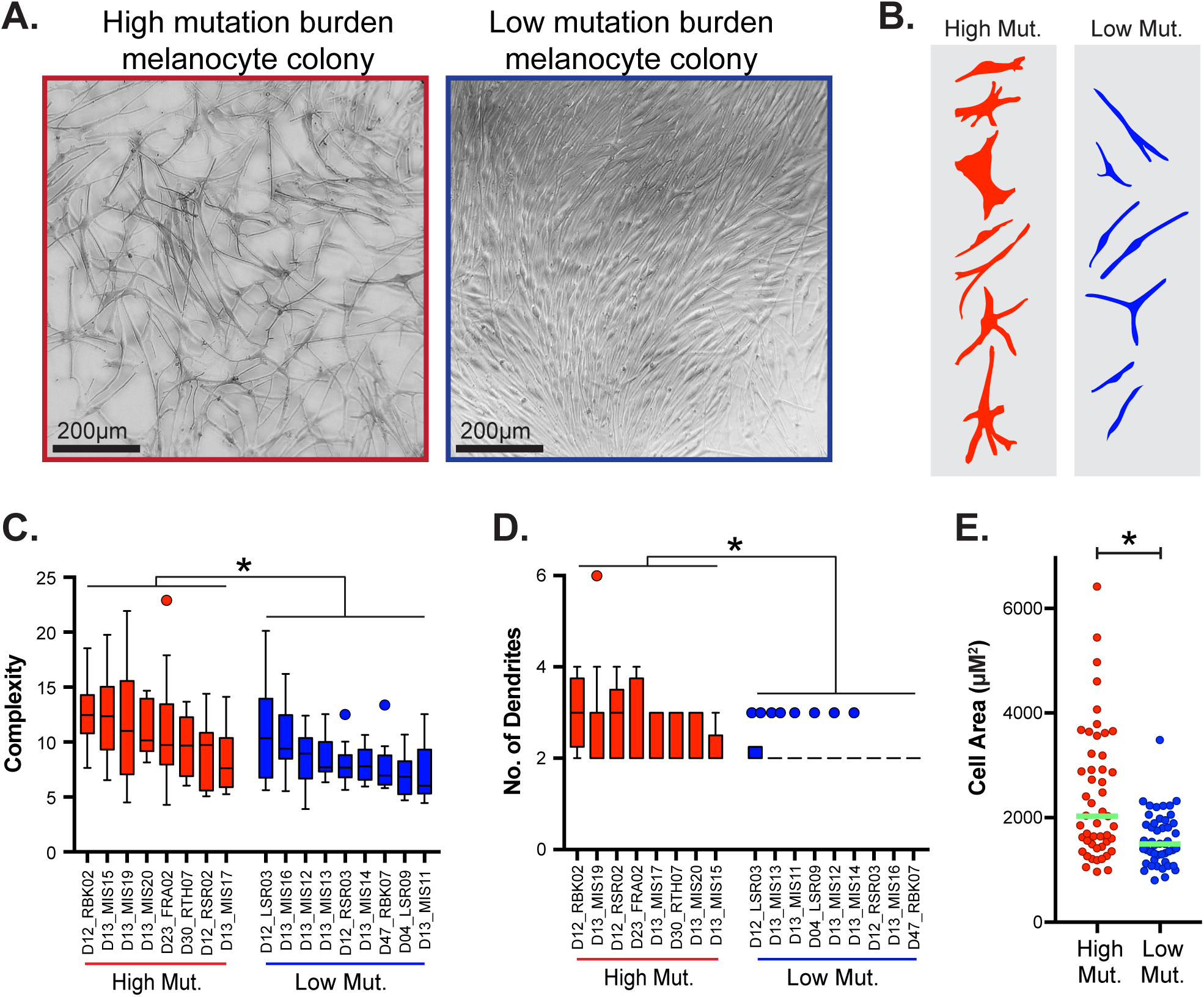
Distinct morphological features in melanocytes with high- and low-mutation burdens. **A.** Phase contrast images of a melanocyte colony with a high mutation burden (Left, 11.27 mut/MB) and a melanocyte colony with a low mutation burden (Right, 2.45 mut/MB). The founding melanocytes for each colony came from the earlobe of Donor 13. **B.** A masked image of individual melanocytes with high vs low mutation burdens, exemplifying the complexity (see methods) of cells with relatively higher mutation burdens. **C**, **D** and **E.** The figures compare cell complexity, dendrite counts and cell area of representative melanocytes from clonal expansions of melanocytes with high and low mutation burdens. Statistical significance was determined using the Student’s t-test (*\*p < 0.005* for comparisons of mean complexity, mean dendrite number and cell area).

Immunostaining for MITF and MLANA (markers for melanocytes) has shown that dendritic melanocytes are enriched in the interfollicular epidermis and the bulb of the hair follicle, while ovoid melanocytes are enriched in the hair bulge^20^ – the niche for epithelial and melanocytic stem cells. Since low mutation burden melanocytes were less dendritic in tissue culture, more stem like, and less sun damaged, we hypothesized that they have spent most of their lives in the hair bulge.

### Spatial transcriptomics reveals the localization of melanocytes with high and low mutation burdens in human skin

To test our hypothesis, we used the Xenium analyzer (10X Genomics) to interrogate the localization of melanocytes with high and low mutation burdens in human skin. The Xenium platform measures spatial expression patterns across hundreds of transcripts at subcellular resolution over a large field of view (up to 1x2cm). When choosing probes, we started with a generic skin panel, designed to mark common cell lineages in the skin, and supplemented it with custom targets, which included all genes upregulated in melanocytes with high and low mutation burdens (see Table S5 for a list of all 360 genes included on the panel). We chose a formalin-fixed paraffin-embedded (FFPE) sample of normal skin from the back of a 63-year-old male who had been diagnosed with melanoma. The skin came from the non-lesional portion (i.e. the normal, adjacent skin) of a wide excision biopsy of the patient’s melanoma. The skin sample also had solar elastosis, suggesting it had experienced high levels of cumulative sun damage. In total, we measured the localization of ∼96 million transcripts in ∼300K cells. Cell segmentation and clustering distinguished different cell types in the skin (Fig. S4), but for the purposes of this study, we focused our analyses on melanocytes.

In each melanocyte, we counted the number of transcripts mapping to genes upregulated in high versus low mutation burden cells and rank ordered cells by their relative expression of these gene sets. Melanocytes with HighMut gene expression programs were almost exclusively found in the interfollicular epidermis (Fig. 5A-C). Melanocytes with LowMut gene expression programs were enriched in hair follicles but could be found in the epidermis, too (Fig. 5A-C). When melanocytes with LowMut gene expression programs were in the epidermis, they were more common near the opening of the hair shaft (Fig. 5B, C). These findings suggest that melanocytes with low mutation burdens originate in the hair follicle but can migrate to the interfollicular epidermis. We also inferred the area of each cell, and the LowMut melanocytes were relatively smaller (Fig. 5D), consistent with the morphological features observed *in vitro* (Fig. 4).

**Figure 5.**
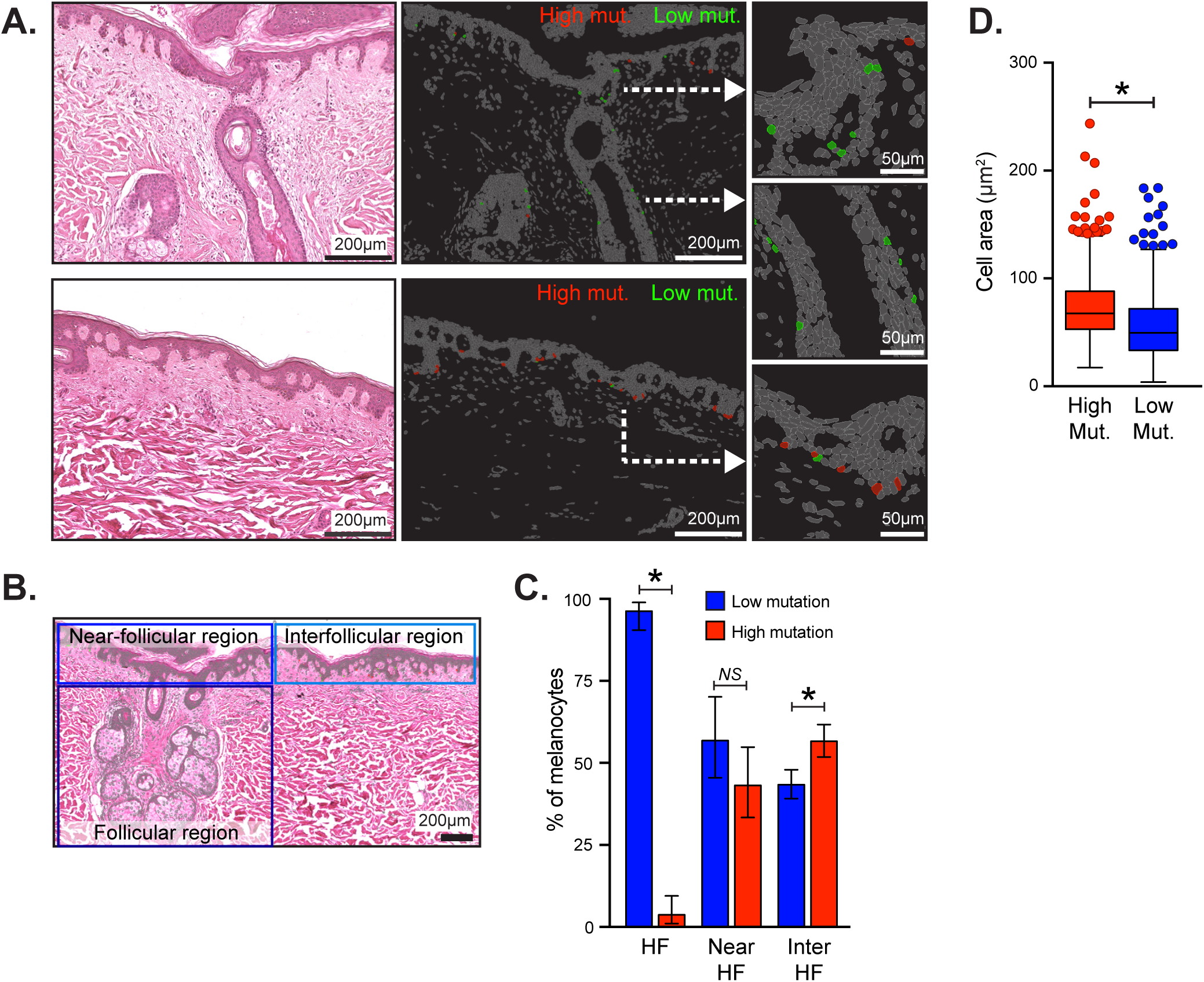
Melanocytes with high- and low-mutation burdens occupy distinct niches in the skin. **A**. Representative examples of a hair follicle (top panels) and interfollicular epidermis (bottom panels). An H&E image of each area is shown on the left, and Xenium cell segmentations are shown on the right with melanocytes colored to indicate which ones are expressing genes associated with high mutation burdens (red, HighMut) versus low mutation burdens (green, LowMut). **B**. LowMut and HighMut melanocytes were counted in the hair follicle (HF), near the hair follicle (Near HF), and between the hair follicles (Inter HF). The hair follicle was almost exclusively populated by LowMut melanocytes. Both HighMut and LowMut melanocytes could be found in the epidermis with LowMut melanocytes more common near the opening of the follicle and HighMut melanocytes more common further away. **C**. The bar graph quantifies the LowMut and the HighMut melanocytes in different skin regions. The error bars show confidence intervals from Exact Poisson test with significant *p*-value (HighMut vs. LowMut) indicated by *. **D**. LowMut melanocytes were smaller in cell area than HighMut melanocytes. The significance (\**p* < 0.001, HighMut vs. LowMut) is calculated by Tukey’s post-hoc HSD test.

## Discussion

In this study, we catalogued mutational, gene expression, morphological, and spatial profiles of melanocytes to infer cellular hierarchies in human skin. Among these modalities, mutational information is the least common in cell atlas studies, but it proved to be uniquely informative. Cells can change their transcriptional states and positional coordinates over time, but mutations are like scars, providing a historical record of the homeostatic processes that were operative on each cell. To our surprise, we found a population of melanocytes in sun-exposed skin with remarkably low mutation burdens, raising an important question. How can cells from the same skin biopsy, ostensibly receiving the same doses of UV radiation, accumulate such different mutation burdens?

Our single-cell analyses were performed on melanocytes harvested from the interfollicular epidermis. A spatial analysis showed that the interfollicular epidermis houses melanocytes inferred to have both high and low mutation burdens, based on their gene expression patterns. These analyses also showed that the hair follicle is heavily enriched with melanocytes inferred to have low mutation burdens, based on their gene expression profiles. We presume that melanocytes with low mutation burdens spend most of their lives in the hair follicle, thus shielding them from the damaging effects of UV radiation, and they migrate to the epidermis in response to UV radiation (Fig. 6). The bulge houses melanocyte stem cells, and the melanocytes with low mutation burdens retain morphological and gene expression features of stem cells.

**Figure 6.**
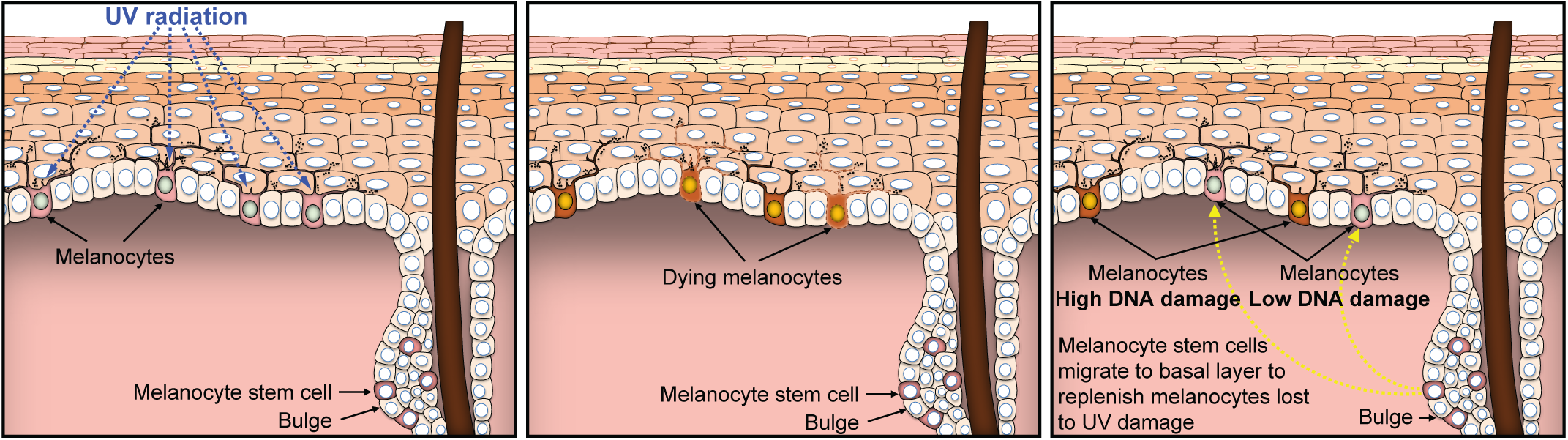
A model to explain how melanocytes with both low- and high-mutation burdens colonize human skin.

In mice, exposure to UV radiation can induce melanocytes to exit the hair follicle and populate the interfollicular epidermis^21^. In humans, patients with vitiligo do not have epidermal melanocytes (a consequence of auto-immunity against those cells), but treatment with UV radiation causes melanocytes in the hair follicles to migrate into the epidermis and repigment the skin^22^. Our work suggests that these waves of migration are not one-off events, only observed in an experimental or disease setting. Instead, they likely occur on a regular basis in physiologically normal skin, perhaps as a mechanism to replenish melanocytes after sun damage (Fig. 6). The possibility that these homeostatic mechanisms could be co-opted for therapeutic intent is intriguing. The mutation burden of tissues is thought to increase with age^23^, but with more research, it may prove feasible to replace heavily mutated cells with fresh cells to rejuvenate tissue.

In a separate study^24^, Yoshida and colleagues described a population of bronchial epithelial cells with “near-normal” mutation burdens admixed with heavily damaged cells from the lungs of smokers. They postulated that the epithelial cells with low mutation burdens may have occupied a privileged niche for much of their lives. In barrier organs, such as skin and lung, more work is needed to understand the location of stem cell reservoirs and how they replenish the cells at the first line of defense. To achieve this goal, cell atlas studies will need to focus on more data modalities than gene expression. Large-scale consortiums have transcriptionally profiled over 85 million cells and counting^9^. While these studies have undoubtably proven useful, our work illustrates the benefits of fewer cells but more information per cell, including mutational profiles.

## Methods

### Data Availability

The genomic and transcriptomic sequencing data for individual cells are available through dbGaP (phs001979.v1.p1 and phs003683.v2.p1). Additionally, the Xenium spatial transcriptomics data for skin sections can also be accessed via GEO (GSE286964). All other results are provided as supplementary materials in the manuscript.

### Skin sample collection for single cell genotyping of melanocytes

A total of 55 skin biopsies were collected from various anatomical sites across 31 donors at the University of California, San Francisco (UCSF) and Northwestern University. Specifically, at UCSF, the biopsies were obtained from cadavers donated through the Willed Body Program for research purposes. At Northwestern, the biopsies were taken from patients recruited at the dermatology clinic. Consent from all living donors was obtained in accordance with protocols approved by the institutional review boards of the respective universities (UCSF IRB 22-36678 and Northwestern IRB STU00211546). For deceased donors, informed consent was part of their will.

The biopsies were collected as either punch or shave biopsies, with diameters of 3mm or 5mm. Additionally, tissue samples (buccal mucosa, blood, or skin biopsies from distinct anatomical sites) were also collected to establish the genome for each donor.

### Genomic and transcriptomic sequencing of individual melanocytes

The skin biopsy was first trimmed to remove excess subcutaneous tissue and then incubated in 10 mg/ml dispase II for 16-18 hours at 4°C. After incubation, the epidermis was separated from the dermis. A single-cell suspension of the epidermal layer was obtained by incubating the tissue in 0.05% trypsin for 3 minutes, with intermittent vortexing every 10-15 seconds.

The resulting single-cell suspension was seeded in CNT40 medium (CELLnTEC) supplemented with 5% antibiotic-antimycotic and cultured at 37°C with 5% CO2 to establish a bulk culture of keratinocytes and melanocytes over the course of a week. Melanocytes were then selectively isolated through limited trypsinization for 3 minutes, which detaches only the melanocytes, leaving the keratinocytes adhered.

The melanocytes were manually seeded as single cells into 96-well plates using serial dilution and cultured in melanocyte media (CNT40) to grow into clonal colonies of approximately 230 cells or more. Manual seeding was preferred over FACS, as it was less abrasive to primary cells and yielded better results. Colonies were monitored daily to ensure that only single-cell-derived melanocyte colonies were selected, and this regular screening also allowed for determining the optimal time to harvest the cells before they underwent growth arrest. Additionally, this approach enabled confirmation of cell type based on cell morphology. The ex vivo clonal expansion increased starting material such that somatic alterations could be detected with high sensitivity and specificity, as previous described^11^ and detailed further below.

The colonies of melanocytes were harvested in RLT buffer (Qiagen, 79216). DNA and mRNA were extracted, separated, and amplified using the G&T sequencing protocol^25,26^. In this method, the DNA was amplified using either multiple displacement amplification (MDA, Qiagen, 150345) or primary template-directed amplification (PTA) protocol^27^ (BioSkryb, 100136). The mRNA was amplified following the SMART-Seq2 protocol as previously described^25,26^. The cDNA amplification was achieved using KAPA HiFi HotStart ReadyMix kit (Roche, KK2502) followed by purification using KAPA HyperPure beads (Roche, 8963843001).

For the reference DNA, bulk DNA was isolated from buccal swabs using the prepIT.L2P method (DNA Genotek, PT-L2P-5) or from unrelated skin biopsies, extracted using the DNeasy Blood & Tissue Kit (Qiagen, 69504).

For library preparation, bulk DNA, melanocyte DNA, or cDNA was sheared using the Covaris LE220 to an average fragment size of 350 base pairs. This was followed by end repair, ligation with IDT8 or IDT10 dual index adaptors, and amplification using the KAPA HyperPrep Kit (Roche, KK8504). DNA libraries were then enriched using the KAPA HyperCapture Reagent Kit (Roche, 09075828001).

During the enrichment/hybridization step, probes were used from either the UCSF500 targeted cancer gene panel (developed and validated by UCSF Clinical Cancer Genomics Laboratory, Roche), NimbleGen SeqCap EZ Exome + UTR panel (Roche, 06740294001), or KAPA HyperExome V1 panel (Roche, 09062556001). The specific exome probe used for each melanocyte is listed in Table S1. Samples were then subjected to paired-end sequencing (100bp) on either an Illumina HiSeq 2500 or NovaSeq 6000, with a total of 297 melanocytes sequenced. For melanocytes with low somatic mutation counts, rehybridization and additional exome sequencing were performed as needed. A graphical summary of this method is shown in figure S1A.

All the nucleic acid inputs and outputs involving G&T-Seq, library preparation, hybridization, and sequencing were quantified using Qubit (dsDNA High Sensitivity quantification), Agilent Bioanalyzer 2100 (High Sensitivity DNA run), and/or QuantStudio 5 real-time PCR system (qPCR with the KAPA Quantification Kit, Roche, KK4854).

### Workflow for sequencing data analysis and variant calling

DNA sequencing data were aligned to the hg19 version of the human genome using the BWA-MEM algorithm (v2.0.5)^28^. Following alignment, Picard (v4.1.2.0) (https://broadinstitute.github.io/picard/) was used to deduplicate the genomic reads. The aligned reads were then realigned around indels and recalibrated using the Genome Analysis Toolkit (GATK v4.1.2.0)^29^. In terms of the RNA samples, sequencing reads were aligned to both the genome and transcriptome using STAR align (v2.1.0)^30^. Deduplication of RNA reads was performed using Picard (v4.1.2.0). Read counts for each gene were quantified with RSEM (v1.2.0)^31^.

To identify germline heterozygous SNPs, FreeBayes (v1.3.1)^32^ was employed, and the SNPs were filtered to include only those overlapping with known SNPs in the 1000 Genomes Project^33^ and with allelic frequencies between 40-60%. Among other uses, this information was utilized to detect allelic dropout, assisting in the removal of samples with low coverage or amplification biases. CNVkit (v0.9.6.2) was used to infer copy number alterations from both DNA and RNA sequencing data^34,35^. Additionally, short insertions and deletions were detected using Pindel^36^, with filtering to retain only indels present in the reference genome and supported by at least four reads. The remaining indels were manually inspected in IGV to remove artifacts.

Somatic mutations were identified as detailed here^11^. Briefly, the initial candidate point mutations were called using MuTect2 (v4.1.2.0)^37^. Although Phi29 polymerase exhibits high fidelity, artifacts can still arise during amplification in MDA or PTA protocols. To remove these and other sequencing artifacts, a multi-step filtering process was applied. First, haplotype phasing was performed using germline heterozygous SNPs to infer haplotype information; true mutations were expected to show linkage to these nearby SNPs, while artifacts typically displayed incomplete or no linkage. Second, RNA validation was conducted by verifying the presence of somatic mutations in RNA sequencing data, as artifacts were not anticipated to be present in both genomic and transcriptomic datasets. This strategy was particularly effective for identifying mutations in highly expressed genes.

Remaining mutations were further scrutinized based on their variant allele frequency (VAF). True mutations exhibited a normal distribution centered around 50%, while artifacts typically had lower allelic frequencies and fewer overall reads. Cutoff thresholds were set using receiver operating characteristic (ROC) curves, trained on known mutations/artifacts, to maximize both sensitivity and specificity.

The inferred mutations (Table S2) are used to calculate the mutation burden as a measure of DNA damage and presented as mutations per megabase. The total megabases of coverage is denoted as footprint of the genome covered. This is calculated using footprint software^11^ and included only nucleotide base pairs with a minimum of 10X or greater coverage.

### Distinguishing melanocyte subpopulations with high and low mutation burden (figure 1)

To identify melanocyte subpopulations with high and low mutation burden, we plotted the mutation burden of individual melanocytes versus the mutation burden of the skin biopsy from which they were derived. For this analysis, the median mutation burden of all sequenced melanocytes from each biopsy was considered the biopsy’s overall mutation burden. Linear regression was performed to fit trendlines through the upper and lower quartiles of mutation burdens. Melanocytes falling into the top quartile (Q1) were classified as having a high mutation burden, while those in the bottom quartile (Q4) were classified as having a low mutation burden (see Fig. 1). Only biopsies with a minimum mutation burden of 3 or higher were included, as it is unreliable to distinguish subpopulations from biopsies with minimal DNA damage.

### Mutation signature analysis between the melanocyte subpopulations (figure 2)

Mutation signatures were inferred for all the cells with a minimum of 10 mutations (Table S2). Cells with fewer than 10 mutations were re-sequenced using probes with broader genome coverage. The mutations from each melanocyte were compiled to create a mutational profile in the trinucleotide context of single base substitutions (SBS), with the Bioconductor library BSgenome.Hsapiens.UCSC.hg19 (v1.4.3) using the DeconstructSigs R package (v1.9.0)^38^. Alexandrov *et al.* previously used non-negative matrix factorization (NMF) to derive 78 unique mutation signatures from 2,780 whole-genome variant calls^15^, which have been extensively validated. These signatures, curated by the Wellcome Sanger Institute, are available in the Catalogue of Somatic Mutations in Cancer (COSMIC) database (https://cancer.sanger.ac.uk/signatures/sbs/).

We utilized the SigProfilerAssignment (v0.1.8), which applies custom forward stagewise algorithm to assign the mutational profile of each cell based on these predefined COSMIC signatures (v3.4)^39^. Figure 2A shows the top 8 mutation signatures observed across all cells, including UV-induced signatures. All remaining mutation signatures present in less than 10% of the cells were grouped as “others.” The overall SBS7 fraction for each melanocyte (Fig. 2D) was calculated by summing all four SBS7 signatures (SBS7a-d).

### Analysis of gene expression profile between the melanocyte subpopulations (figure 3, S2)

Differential gene expression analysis was conducted using the DESeq2 R package (v1.38.3)^40^. RNA reads from the RSEM file were first normalized and transformed using variance-stabilizing transformation. A significance threshold of 10% adjusted p-value (Benjamini-Hochberg) and a log2(fold change) greater than 0 was applied to identify differentially expressed genes. To visualize these genes, a volcano plot (Fig. 3A) and a heatmap (Fig. 3B) were generated. Additionally, gene expression in the two melanocyte subpopulations was compared to that of keratinocytes and fibroblasts^41^ using cell type-specific markers: PMEL, TYRP1, and MLANA for melanocytes; KRT14, KRT17, and KRT5 for keratinocytes; and LUM, COL1A1, and S100A4 for fibroblasts (Fig. 3B), to confirm cell identity.

Genes enriched in melanocytes with high and low mutation burdens, as identified through differential expression analysis, were compared to enriched genes from three additional datasets: (a) adult melanocytes vs. melanocyte stem cells (MSC), (b) volar vs. non-volar melanocytes, and (c) foreskin vs. trunk melanocytes. Datasets (a) and (b) were obtained from Belote *et al.*^6^, while dataset (c) was from Cheng *et al*.^18^ The hypergeometric test was performed on overlapping genes (Table S3) in R using the phyper function as follows:

$ phyper(o, a, b, x, lower.tail = FALSE, log.p = FALSE)

Here, “o” represents the number of overlapping genes, “a” is the number of enriched genes in the first set, “b” is the number in the second set, and “x” is the number of remaining genes from the total platform of 18,000 genes. Statistical significance was based on the upper tail of the distribution (lower.tail = FALSE). The overlaps were shown as Venn diagrams (Fig. 3D, S2A and S2B).

We further analyzed the overlap between the enriched gene sets from (a) high mutation burden melanocytes and (b) low mutation burden melanocytes with the annotated gene sets in the Molecular Signatures Database (MSigDB) using the Gene Set Enrichment Analysis (GSEA)^16^ tool. Specifically, we compared these overlaps with the following annotated human gene sets: H (hallmark gene sets), C1 (positional gene sets), C2 (curated gene sets), C3 (regulatory target gene sets), C4 (computational gene sets), C5 (ontology gene sets), C6 (oncogenic signature gene sets), C7 (immunologic signature gene sets), and C8 (cell type signature gene sets)^16,42^. Overall, our gene sets were compared to 33,591 MSigDB gene sets, comprising 42,499 genes. The results are presented as the number of overlapping genes per gene set, with statistical significance indicated by p-values and q-values (false discovery rate). All gene sets that showed significant overlap with either the high- or low-mutation burden melanocyte subsets are listed in Table S4.

### Comparing cell morphology between the melanocyte subsets (figure 4)

Cell images were captured using either the Zeiss Axiovert 40 CFL or the Invitrogen EVOS FL system. Grayscale images were extracted with FIJI, and brightness and contrast were adjusted to enhance cell visibility. The “rolling ball” algorithm-based background subtraction was used to correct unevenly illuminated background.

### Cell segmentation and analysis (figure 4)

Raw in-focus images were processed by researchers blinded to mutational status. Briefly, brightfield images were imported into QuPath. The cells that could be distinguished from their neighbors were manually segmented using the polygon tool (4-15 cells per field of view). Annotation measurements were exported, and complexity was calculated from a perimeter-to-area ratio following the method from Hou 2018^43^. Additionally, the number of dendrites, or projections from the cell body, were manually counted for each segmented object. Measurements were analyzed in GraphPad Prism. A representative selection of ROI’s was exported to Fiji and arranged using the ROI manager. The masks of the cells were filled and sorted based on high and low mutation. The complexity and dendrite count of representative melanocytes from clonal expansions of melanocytes with high and low mutation burdens were visualized using in a box and whisker plot using Tukey’s method, with significance calculated by Student’s t-test (Fig 4C, D).

### Spatial transcriptomic profiling using Xenium platform

#### Sample selection and sectioning

For the spatial transcriptomics analysis of melanocyte subsets with high and low mutation burdens using the 10X Xenium platform, we obtained a formalin-fixed paraffin-embedded (FFPE) skin sample from a 63-year-old male donor. The skin mount consisted of normal, non-cancerous tissue located at the margin of a wide excision performed to remove melanoma, ensuring the presence of normal melanocytes with a high mutation burden. A 5 µm section of the skin was mounted on a Xenium slide, with two skin sections positioned within the sample area (10.45 mm x 22.45 mm) such that the epidermis of each section faced the other along the length of the slide. Care was taken to ensure that none of the fiducial markers were covered, and any obstructions were manually removed to expose all markers correctly. Two adjacent sections were mounted on two separate Xenium slides for dual Xenium analysis.

Prior to mounting on the Xenium slides, some sections were first mounted on standard slides and stained with H&E to verify the integrity of the tissue. Additionally, sections were subjected to high-sensitivity eukaryotic Pico RNA analysis (Agilent, 5067-1513) using an Agilent Bioanalyzer 2100. The DV200 score was confirmed to be within the acceptable range for the Xenium run (50-70%).

### Gene panel design

The targeted gene panel was based on the predesigned human skin panel (v1), which comprised of 260 genes. An additional 100 genes were incorporated to create a custom panel. These additional genes cover various melanocyte subtypes, including those enriched in high-mutation melanocytes (n = 20), low-mutation melanocytes (n = 15), adult melanocytes (from Belote *et al.*; n = 12), neonatal melanocytes (from Belote *et al.*; n = 9), fetal melanocytes (from Belote *et al.*; n = 3), melanocyte stem cells (from Belote *et al.* and other reports; n = 12), differentiated melanocytes (n = 6), transitional melanocytes (n = 3), neural crest cells (n = 17)^44–56^, hair follicle bulge cells (n = 2)^57,58^, and Schwann cells (n = 1)^59^. Of the 23 genes enriched in high-mutation melanocytes and 19 genes in low-mutation melanocytes, some of them were already included in the predesigned skin panel. The genes FRG2DP and FTH1P10 were excluded due to the lack of suitable probes. The genes are listed in Table S5.

Although most custom-added genes had the maximum probe set coverage (up to 8 probes), some were reduced to prevent optical crowding. Additionally, the panel includes 40 negative control codewords to assess the specificity of the decoding algorithm and 20 negative control targets to evaluate the assay’s specificity.

### Sample preparation for Xenium assay

The slides were processed according to the 10X Genomics recommended protocol. Both slides were handled concurrently using the Xenium Slides & Sample Prep Reagents (10X Genomics, PN-1000460). First, the slides were assembled into cassettes (10X Genomics, PN-1000566). Next, the paraffin was removed from the slides, followed by rehydration and decrosslinking of the tissue sections to release sequestered mRNA.

Next, probes were hybridized to the target mRNA in the skin sections at 50°C for 16-24 hours. After hybridization, excess probes were washed off during a post-hybridization wash for 30 minutes at 37°C. The probes consist of three regions: two regions at the ends that hybridize to the target RNA, and a middle region that encodes the gene-specific barcode.

Ligation of the probes to the target RNA forms a circular DNA probe structure. This is accomplished with a 2 hour incubation at 37°C. Following this, rolling circle amplification is performed for 2 hours at 30°C, generating multiple copies of the gene-specific barcode for each RNA target.

### Cell segmentation staining

To accurately identify the borders of each cell, immunofluorescence staining was performed using various cell structure markers, including boundary stains (TP1A1, CD45, E-Cadherin) and an interior RNA stain (18S). The staining was carried out by incubating the slides at 4°C for 16-24 hours using the Xenium Cell Segmentation Staining Reagents (10X Genomics, PN-1000661). After incubation, the stain was enhanced by adding the Xenium Staining Enhancer reagent, followed by washing to remove excess staining reagents. Autofluorescence was then quenched to reduce background noise. The final step in the staining process involved nuclear staining with DAPI.

### Xenium Onboard Analysis (XOA)

The slides were processed using the fully automated Xenium Analyzer. The skin samples were imaged over 15 cycles, where fluorescent probes targeting mRNA sequences and other reagents were added, imaged, and removed during each cycle. The image sensor captured 3D data across four fluorescence channels and multiple fields of view (FOV) throughout these cycles. The resulting images were stitched together to create a comprehensive spatial dataset for Xenium analysis.

The Xenium Analyzer then processed and decoded the transcript data using its onboard probabilistic model through the XOA pipeline (v3.0.2.0). This decoding utilized predefined fluorescent signal patterns, or codewords, to identify transcripts, with negative codewords included to ensure specificity. Once decoded, the transcripts were deduplicated, and the final data underwent secondary analysis to produce the XOA output. The analysis summary provided information on decoding accuracy, cell segmentation, gene expression and image quality control (QC). All the outputs were visualized using Xenium Explorer (3.1.0).

### Post Xenium H&E staining of skin sample

Taking advantage of the non-destructive Xenium workflow, the skin sample slides were H&E-stained post analysis run. During the cell segmentation staining, the autofluorescence quenching leaves behind a purple stain. Therefore, the slides were first removed from their cassettes, and quencher removal was performed using 10 mM sodium hydrosulfite (Sigma-Aldrich, 157953-5G). The skin sections were then stained with Hematoxylin and Eosin, dehydrated, and coverslipped for imaging. The samples were scanned at 20X magnification using the Leica Aperio CS2. The output was processed using QuPath bioimage analysis software to extract a pyramidal, tiled “OME.TIF” image file. This image was then imported into Xenium Explorer and manually aligned using a best similarity transformation method. The alignment relies on key points identified from both the XOA spatial output and the H&E-stained image.

### Spatial analysis of high- and low-mutation melanocytes

Decoding for two skin samples yielded 84.4% (48,418,166) and 84.9% (46,940,134) high-quality gene transcripts, with cell segmentation detecting 148,741 and 145,489 cells, respectively. In the first sample, 28.2% of the cells were segmented based on boundary staining, 62.8% by RNA staining, and 9.0% by nuclear (DAPI) expansion. The second sample showed similar segmentation, with 27.5% boundary stain, 62.5% RNA stain, and 10.0% DAPI expansion. For further analysis, cells with fewer than 20 transcripts per cell, as determined by the Phred quality score, were excluded. The XOA pipeline identified 41 distinct clusters in the first skin sample based on gene expression profiles. These clusters were categorized into five groups: melanocytes, keratinocytes, immune cells, adnexal cells, fibroblasts, and an “others” category, which likely included apoptotic cells, unevenly segmented cells, or rare cell types that do not fit into the other five main categories. The reannotated clusters were visualized using a UMAP projection (Fig. S4). A total of 1,918 melanocytes were detected.

To validate these annotations, the Xenium output was reanalyzed using the Seurat R toolkit (v5.1.0)^60^. Multi-cluster differential expression analysis using the DESeq2 package (v1.38.3) confirmed the defined cell types (Fig. S4). The thresholds for this analysis were an adjusted p-value (Benjamini-Hochberg) of 0.05 and a log2 fold change greater than 1. The top 10 genes enriched in each cell type cluster were visualized as a dotplot heatmap, where upregulated genes were marked in red, and the size of the dot indicated the fraction of cells expressing the gene.

To identify melanocytes with high and low mutation burdens, a read count matrix for all genes in the cluster was generated. A delta count (the difference between the sum of reads for genes enriched in high mutation melanocytes and those enriched in low mutation melanocytes) was calculated for each cell, and cells were rank ordered based on their relative expression of “HighMut” versus “LowMut” genes. The top 15th percentile of cells was classified as high mutation melanocytes, and the bottom 15th percentile as low mutation melanocytes. This method relies on detecting cells with high expression of genes associated with high mutation melanocytes and low expression of genes associated with low mutation melanocytes, and vice versa. Using this approach, 287 high mutation melanocytes and 287 low mutation melanocytes were identified in the first sample, and 307 high mutation melanocytes and 307 low mutation melanocytes in the second sample.

After identifying the two melanocyte subpopulations spatially, we quantified their counts in three distinct regions of the skin sample based on proximity to hair follicles: follicular, near-follicle, and interfollicular regions (Fig. 5). The results are presented as a bar graph with a 95% confidence interval, and significance was determined using the Poisson test. We also measured the surface area of each melanocyte in the two subpopulations. The surface area data was visualized in a box and whisker plot using Tukey’s method, with significance calculated using Tukey’s post-hoc Honest Significant Difference (HSD) test.

## Acknowledgements

The study was supported by grants from: NIH NIAMS (AR080626), NIH NCI Human Tumor Atlas (HTAN) network (U01 CA294536), NIH NCI (R01 CA265786), Department of Defense Melanoma Research Program (ME210014), Melanoma Research Alliance (Dermatology Fellows Award), the LEO Foundation Region Americas Award, UROP from the Office of Undergraduate Research at the University of Utah (awarded to Kayla Marks), and the UCSF Department of Dermatology.

---

**Figure S1.**
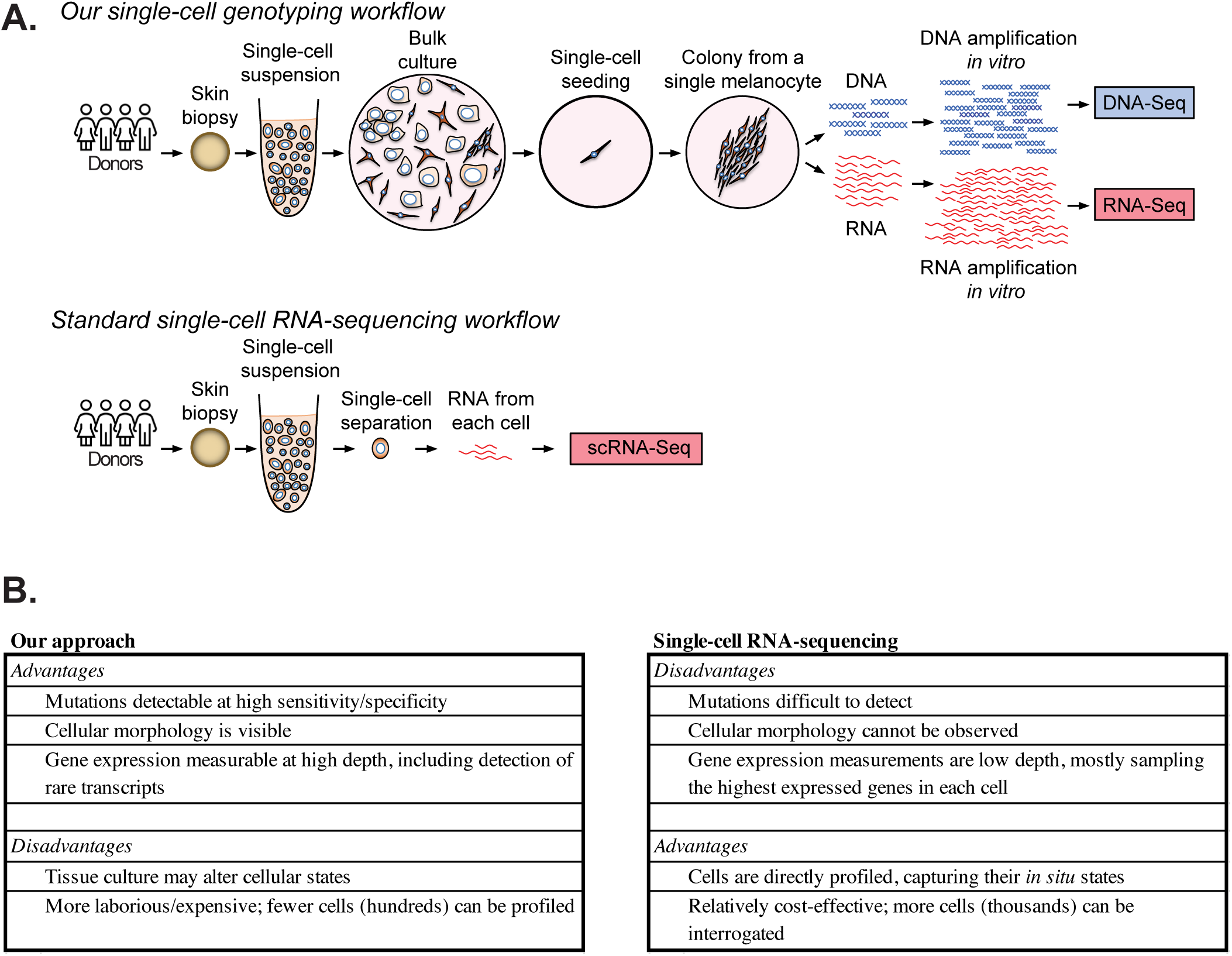
An approach to measure mutational, gene expression, and morphological features from small expansions of individual melanocytes. **A.** An overview of our single-cell genotyping workflow compared to a typical single-cell RNA-sequencing workflow. **B.** Advantages and disadvantages of our workflow compared to a standard single cell RNA sequencing workflow.

**Figure S2.**
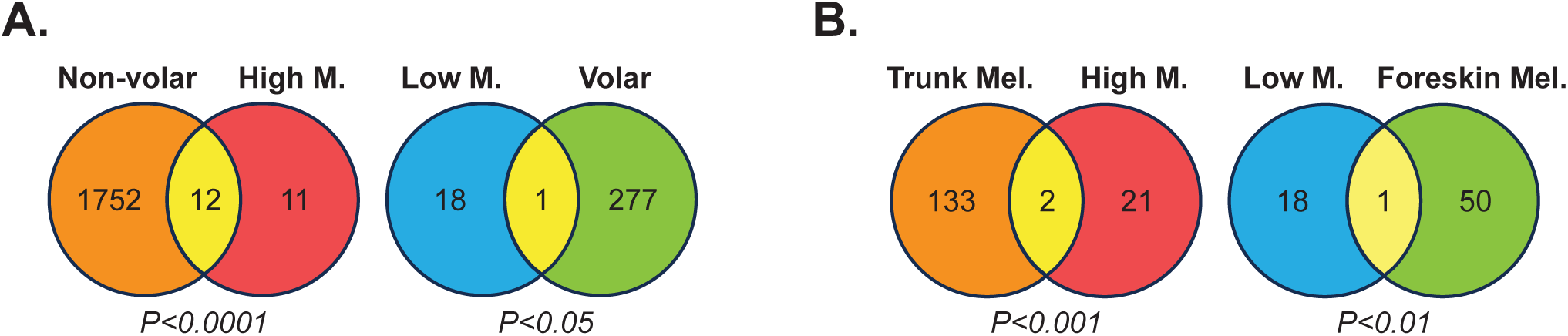
Genes expressed in high- and low-mutation melanocytes show contrasting overlap with gene lists from other studies and contrasting morphology. The Venn diagrams show the overlap of expressed genes (see figure 3) in melanocytes with high and low mutation burden with: **A.** genes preferentially expressed in non-volar or volar melanocytes (from Belote *et al.* Nature Cell Biology, 2021) and **B.** differentially expressed genes from trunk or foreskin melanocytes (from Cheng *et al.* Cell Reports, 2018). The statistical significance of overlap is calculated using a hypergeometric distribution.

**Figure S3.**
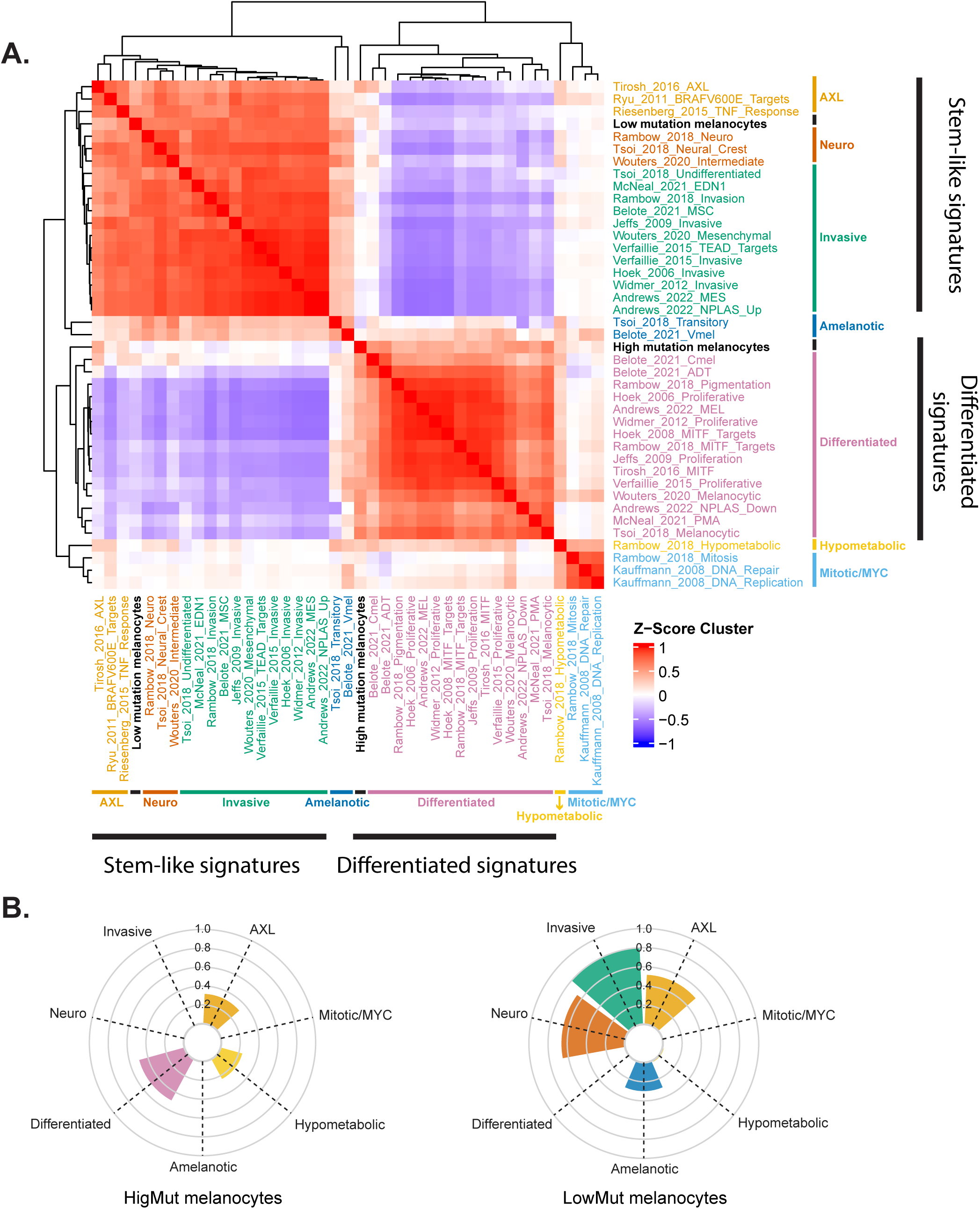
The relationship between HighMut and LowMut gene expression signatures relate to other melanocytic signatures. The WIMMS (What Is My Melanocytic Signature) tool was used to determine how gene expression in high- and low-mutation burden melanocytes correlate with genes expressed in previously published melanocytic signatures. Previously published signatures are grouped broadly into stem-like and differentiated signatures with subgroups, as indicated. Our HighMut signature clustered with other differentiated signatures while our LowMut signature clustered with other stem-like signatures.

**Figure S4.**
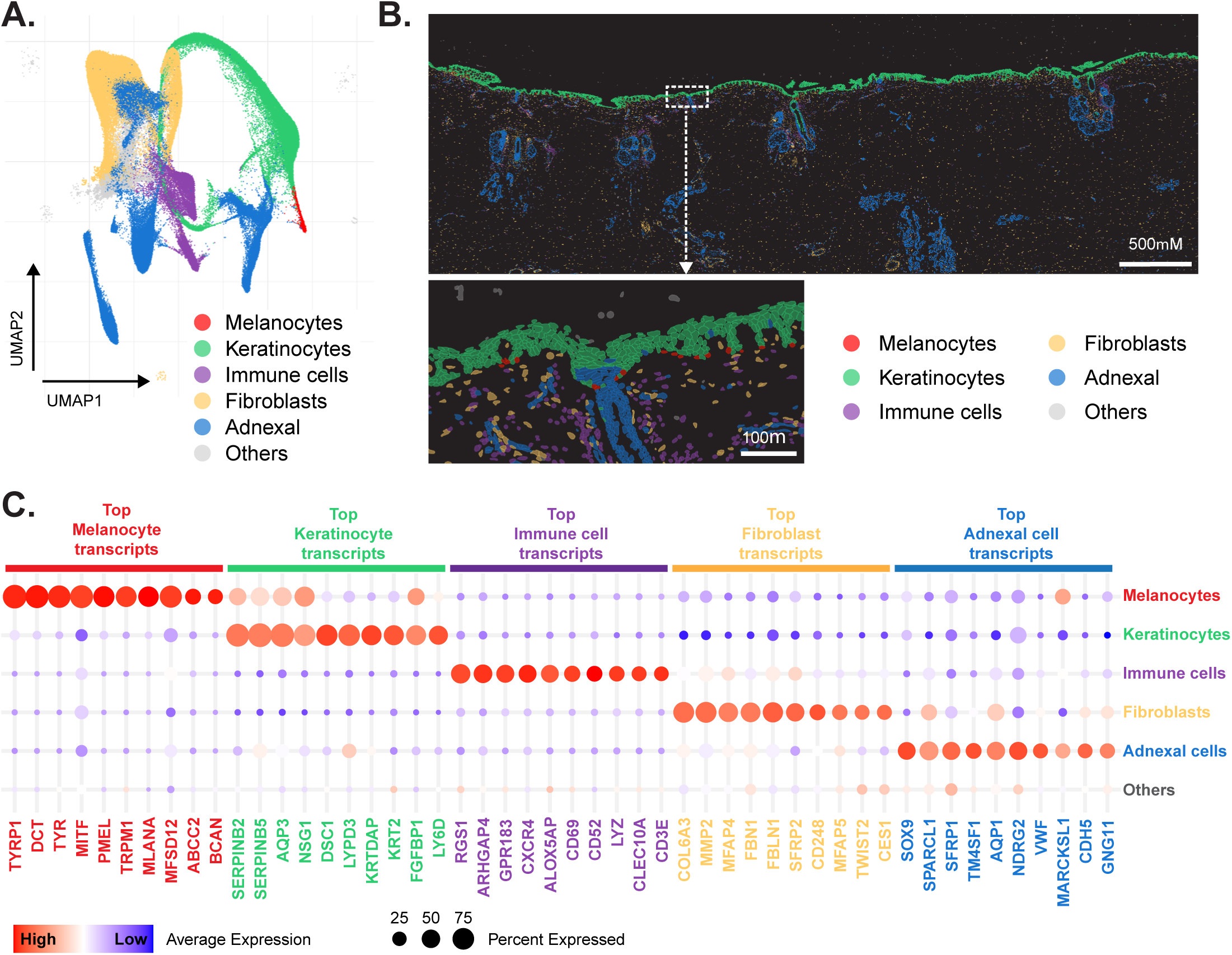
Diverse cell types, including melanocytes, are recognized in Xenium data of normal human skin. **A.** A UMAP, produced by the Xenium Ranger workflow (see methods), shows the relationship of hundreds of thousands of cells from a normal skin biopsy. The graph-based clustering algorithm identified 42 clusters of cells, which were aggregated into six broader groups of cells based on their gene expression profiles. **B.** The localization of different cell types in Xenium data. The epidermis is mainly comprised of keratinocytes with melanocytes occasionally observed in the basal layer. Adnexal structures extend into the dermis, surrounded by a low density of fibroblasts and immune cells. **C.** The top 10 genes associated with each group of cells. The melanocytes were distinguished by their expression of known pigmentation genes.

